# *CircSry* regulates spermatogenesis by enhancing γH2AX expression via sponging miR-138-5p

**DOI:** 10.1101/2021.10.25.465684

**Authors:** Yanze Song, Min Chen, Yingfan Zhang, Na Li, Min Chen, Miaomiao Qiao, Yuanwei Cao, Jian Chen, Fei Gao, Haoyi Wang

## Abstract

*Sry* on the Y chromosome is the master switch in sex determination in mammals. It has been well established that *Sry* encodes a transcription factor that is transiently expressed in somatic cells of male gonad, inducing a series of events that lead to the formation of testes. In the testis of adult mice, *Sry* is expressed as a circular RNA (circRNA) transcript, a type of noncoding RNA that forms a covalently linked continuous loop. However, the physiological function of this *Sry* circRNA (*circSry*) remains unknown since its discovery in 1993. Here we show that *circSry* is mainly expressed in the spermatocytes, but not in mature sperms and Sertoli cells. Loss of *circSry* led to the reduction of sperm number and the defect of germ cell development. The expression of γH2AX was decreased and failure of XY body formation was noted in *circSry* KO germ cells. Further study demonstrates that *circSry* regulates H2AX mRNA indirectly in pachytene spermatocytes through sponging miR-138-5p. Our study demonstrates that, in addition to its well-known sex-determination function, *Sry* also plays important role in spermatogenesis as a circRNA.

## Introduction

Long non-coding RNAs are relatively abundant in the mammalian transcriptome (1), and play important roles in gene regulation in development and reproduction (2). Circular RNA (circRNA) is a unique type of non-coding RNAs generated through back-splicing to form a covalently linked loop (3, 4). Since the first circular RNA was discovered in 1970s (5), very few circRNAs had been identified in the following years. In the last decade, however, the development of RNA sequencing technologies and bioinformatics has greatly facilitated the discovery of circular RNAs (6). Many circular RNAs were found stably expressed in various cell types, and engaged in regulating various biological processes, such as transcription, alternative splicing, chromatin looping, and post-transcriptional regulation (7–12). One of the functional mechanisms of circular RNA is that they act as competing endogenous RNAs (ceRNAs) to sponge miRNAs, therefore regulating gene expression (4, 13–15).

*Sry* is best known as the sex-determination gene on Y chromosome. In mouse, *Sry* is expressed as a transcription factor from 10.5 to 12.5 post-coitum (dpc) in the genital ridge somatic cells, initiating testis development (16). Introduction of *Sry* locus into the female mouse embryo switches the sex to male, while the targeted mutation in male embryos leads to complete male-to-female sex reversal (17–19). Moreover, mutation of *SRY* causes a range of sex-disorder development with profound effects in human (20). Recently, a cryptic second exon of mouse *Sry* hidden in the palindromic sequence was identified and this two-exon *Sry* transcript plays primary role in sex determination (21). *Sry* is also expressed in adult mouse testis as a circular RNA (*circSry*) (22–24). A linear transcript containing long inverted repeats is transcribed from a distal promoter, followed by a back splicing event that covalently links an acceptor splice site at the 5’ end to a donor site at a downstream 3’ end (Figure 1A) (4). Although the presence of *circSry* in the testis has long been discovered, the significance of *circSry* remains elusive.

**Figure 1.**
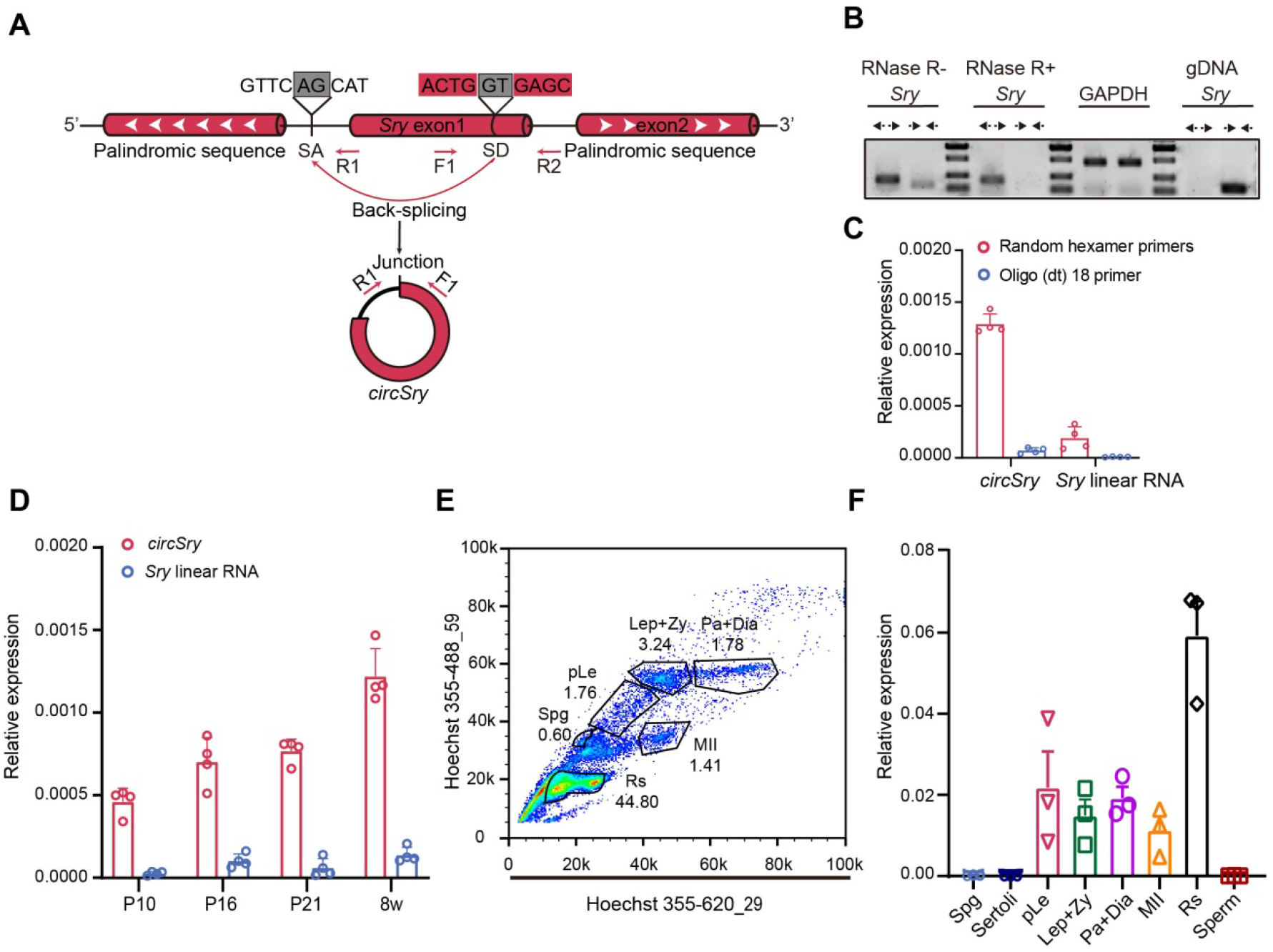
Identification of *circSry*. (A) Scheme illustrating of the generation of *circSry. CircSry* is formed by an incomplete single exon *Sry* gene through back splicing mechanism. Convergent or divergent primers detect circRNA (F1 and R1) or linear RNA (F1 and R2) of *Sry*. Gray box indicates the head-to-tail splicing sequence. (B) Production of divergent primers was resistant to RNase R treatment. *CircSry* was not amplified using genomic DNA as template (n=3); GAPDH was used as control. (C) Random hexamer primer or Oligo (dt)_18_ primers were used to analyze the expression levels of *circSry* in 8 week-old control mice (n=4). (D) Relative expression of *circSry* or linear RNA in testis from 10 days postnatal to adulthood (n=4). (E) Fluorescence cytometry separated different subtypes of germ cells in adult testis and (F) relative expression of *circSry* in 8 types of cells. (Spg: spermatogonium; pLe: pre-leptotene stage; Lep+Zy: leptotene stage and zygotene stage; Pa+Di: pachytene stage and diplotene stage; MII: meiosis II stage; Rs: round spermatids; Sperm: mature sperm; Sertoli: Sertoli cells; n=3). The data are presented as the mean ± s.e.m. Source data is available as a Source Data file.

In this study, we generated mouse models that did not express *circSry*, without interfering with male sex determination. We found that *circSry* played an important role in spermatogenesis, and further dissected the underlying mechanism. Our findings highlight a unique synergy between *Sry*’s male sex-determination role as a protein and its regulatory role as a circular RNA in male germ cells.

## Results

### Characterization of *circSry* in mouse testis

To identify *Sry* transcripts in adult mouse testis, divergent primers and convergent primers were designed to amplify *circSry* or *Sry* linear RNA respectively (Figure 1A). Upon RNase R treatment, *circSry* was still detectable by RT-PCR, while linear RNA was not (Figure 1B). The expression of *circSry* was abundant when random hexamer primers were used for reverse transcription, while only weak signal was detected using oligo (dt)_18_ primers. In comparison, the expression of *Sry* linear RNA was barely detectable using either random hexamer primers or oligo (dt)_18_ primers (Figure 1C). The presence of head-to-tail splicing site of *circSry* was verified by Sanger sequencing (Figure1 figure supplement 1A). Furthermore, by separating the cytoplasm and nucleus fractions, we found that *circSry* was mainly localized in the cytoplasm (Figure figure supplement 1B). All these results confirmed previous reports that *Sry* transcripts in adult mouse testis are non-polyadenylated circular RNAs mainly localized in cytoplasm (24).

Next, we measured the level of *Sry* transcription from day P8 to adulthood. The amount of *circSry* increased over time while linear RNA was barely detectable (Figure 1D). To examine the expression of *circSry* in different cell types of adult testes, different types of germ cells were isolated by flow cytometry (39): spermatogonia (Spg), Pre-leptotene spermatocytes (pLE), leptotene and zygotene spermatocytes (Lep+Zy), pachytene and diplotene (Pa+Di) spermatocytes and round spermatids (Figure 1E). Mature sperms were obtained from adult mouse epididymis and Sertoli cells were obtained from P20 testis. *CircSry* was detected in meiotic and post-meiotic germ cells, expressed at the highest level in round spermatids. There was no *circSry* detected in spermatogonia, mature sperms or Sertoli cells (Figure 1F).

### Generation of *circSry* KO mouse

To determine whether *Sry* is involved in spermatogenesis, we generated *circSry* KO mouse via CRISPR/Cas9 (Figure 2A). We designed a sgRNA to specifically target splice acceptor site upstream of *Sry* coding region. Deletion of the 11 bp region harboring splice acceptor site led to complete loss of *circSry* (Figure 2, B and C). No significant difference was detected in control and *circSry* KO embryo gonads (Figure 2 figure supplement 1A-C). No gross abnormalities of external genitalia were observed in 8-week-old *circSry* KO (XY) male founder mice (Figure 2 figure supplement 2A), and they were fertile. In F0 and F1 generation, all the male mice carried Y chromosome and all the females did not, these results demonstrated that this deletion of 11bp upstream region of *Sry* did not interfere with sex determination (Figure 2 figure supplement 2B). Female offsprings were normal, and no developmental defects were observed (Figure 2 figure supplement 2A).

**Figure 2.**
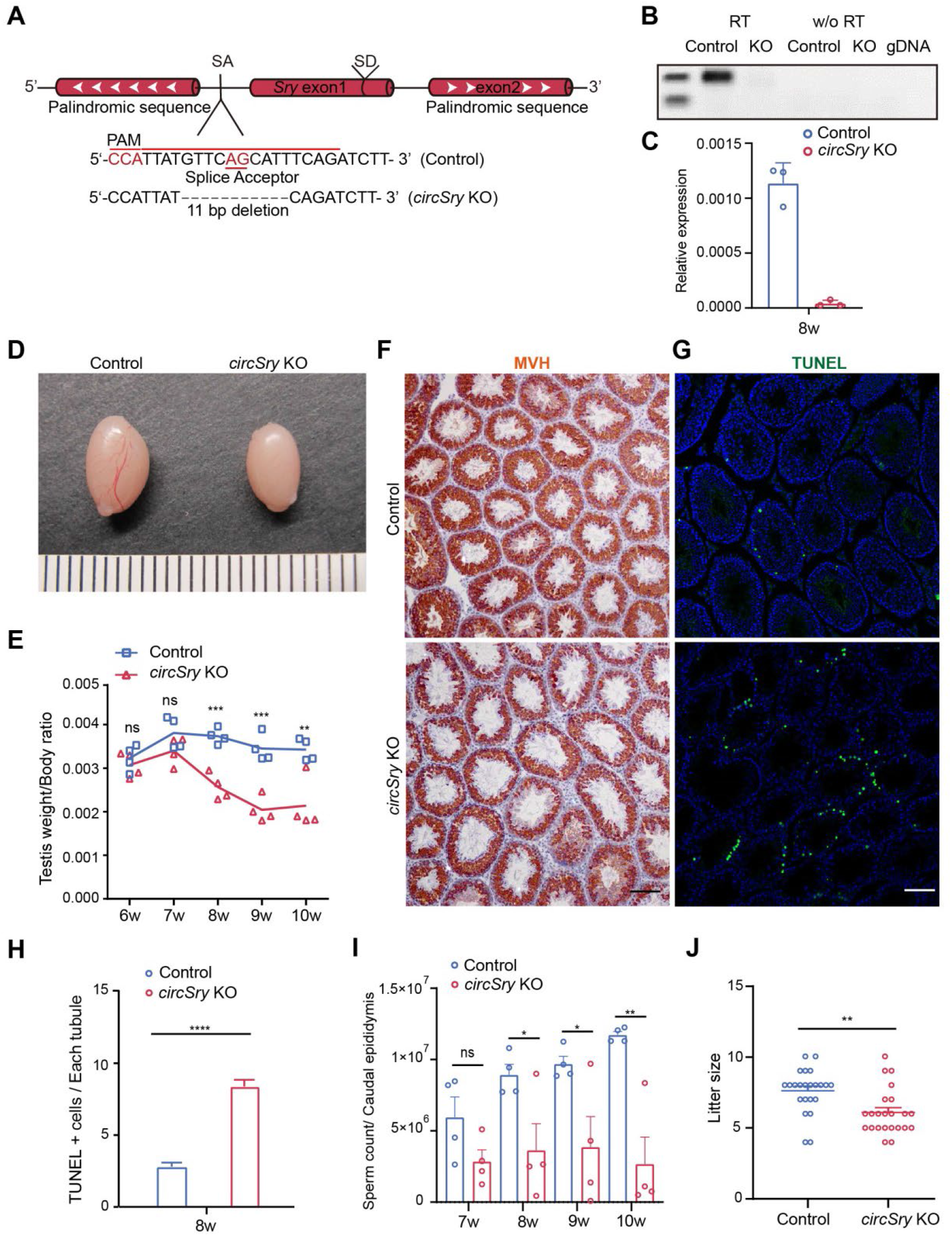
Loss of *circSry* impairs male mice fertility. (A) Design of deleting *circSry* using CRISPR/Cas9; a specific sgRNA was designed to target splicing acceptor site of *circSry*. (B) RT-PCR results of *circSry*. (C) Quatification of *circSry* within control or *circSry* KO mice testes.(D) Representative image of *circSry* KO and control (wild-type) testis of 8-week-old mice. (E) Testis/body ratio of *circSry* KO and control mice from 6 to 10 weeks of age. P values are presented above the relevant bars (*p<0.05, **p<0.01, ***p<0.001, ns, not significant; unpaired, two tailed t test, n=4). (F) Germ cells were labeled with antibody against MVH (brown). Loss of epithelium within the seminiferous tubules in *circSry* KO compared with control mice. Scale bars indicate 100 μm. (G), (H) TUNEL signal in *circSry* KO and control mouse testis. The total tubule number reached 200 (****p<0.0001; unpaired, two tailed t test). (I) Sperm count of *circSry* KO mice and control mice from 7 to 10 weeks of age (*p<0.05, **p<0.01, ns, not significant; unpaired, two tailed t test n=5). (J) Litter size of *circSry* KO compared with control mice. Random 8-week-old males of control and *circSry* KO were chosen to breed with 6-week-old female of C57BL/6 for 4 months (**p<0.01, unpaired, two tailed t test; n=4). Scale bars indicate 100 μm. The data are presented as the mean ± s.e.m.

### The fertility of *circSry* KO male mice was affected

To determine the function of *circSry*, we assessed the testes of 6 to 10 week-old males and examined the germ cell development in *circSry* KO mice. The size of testes from *circSry* KO mice was smaller than that of control (wild-type C57BL/6) mice at 8 weeks (Figure 2D). The testis to body weight ratio of *circSry* KO mice was comparable to that of control mice at 7 weeks, whereas it became significantly lower from 8 to 10 weeks (Figure 2E). The development of germ cells was examined by MVH staining. As shown in Figure 2F, the histology of the seminiferous tubules was grossly normal. MVH-positive germ cells were detected in the testes of both control and *circSry* KO mice at 8 weeks (Figure 2F). However, the lumen size of seminiferous tubules in *circSry* KO mice was larger compared with that of control mice (Figure 2F). There were notably more TUNEL-positive apoptotic cells in the seminiferous tubules of *circSry* KO mice than that of control mice (Figure 2, G and H). Furthermore, we found that the total number of sperms in the caudal epididymis of *circSry* KO mice was lower than that in control mice (Figure 2I). To test the fertility of *circSry* KO mice, we crossed 2-3 month-old *circSry* KO male with wild-type female mice and counted the litter size born within 4 months. *CircSry* KO mice produced smaller litter size than age-matched control male mice (Figure 2J). Notably, no difference of SOX9 positive Sertoli cells was detected between control and *circSry* KO seminiferous tubules (Figure 2 figure supplement 3, A and B), suggesting that Sertoli cells were not affected. In addition, the percentage of mobile sperm remained no difference between the control and the *circSry* KO mice (Figure 2 figure supplement 3C), indicating that the loss of *circSry* did not affect sperm mobility. Taken together, these results suggested that the germ cell development was abnormal in *circSry* KO mice.

### Germ cell specific knockout of *circSry* led to the defects of spermatogenesis

To characterize the cell type specific function of *circSry* in spermatogenesis more rigorously, we generated a *Sry* conditional KO mouse model *Sry^flox^* by inserting two loxP sites flanking *Sry* (Figure 3 figure supplement 1A). *CircSry* was specifically knocked out in germ cells or Sertoli cells by crossing *Sry^flox^* male mice with *Stra8-Cre* or *Amh-Cre* transgenic mice respectively (Figure 3 figure supplement 1A). Compared with control mice, the testes size and the testis to body weight ratio of *Sry^flox^*; *Stra8-Cre* male was smaller at 2 months of age, and the number of sperms in the caudal epididymis was reduced (Figure 3, A and B). Accordingly, decreased MVH-positive germ cells and increased number of apoptotic germ cells were observed in *Sry^flox^*; *Stra8-Cre* mice (Figure 3, E and F, Figure 3 figure supplement 1B). These defects were similar to that observed in the *circSry* KO mice. By contrast, no defect of germ cell development was observed in *Sry^flox^*; *Amh-Cre* male mice (Figure 3, C and D, G and H, Figure 3 figure supplement 1C), indicating that *Sry* is not required for Sertoli cells function in adult male testis. Taken together, we conclude that *circSry* plays an important role in germ cell development during spermatogenesis.

**Figure 3.**
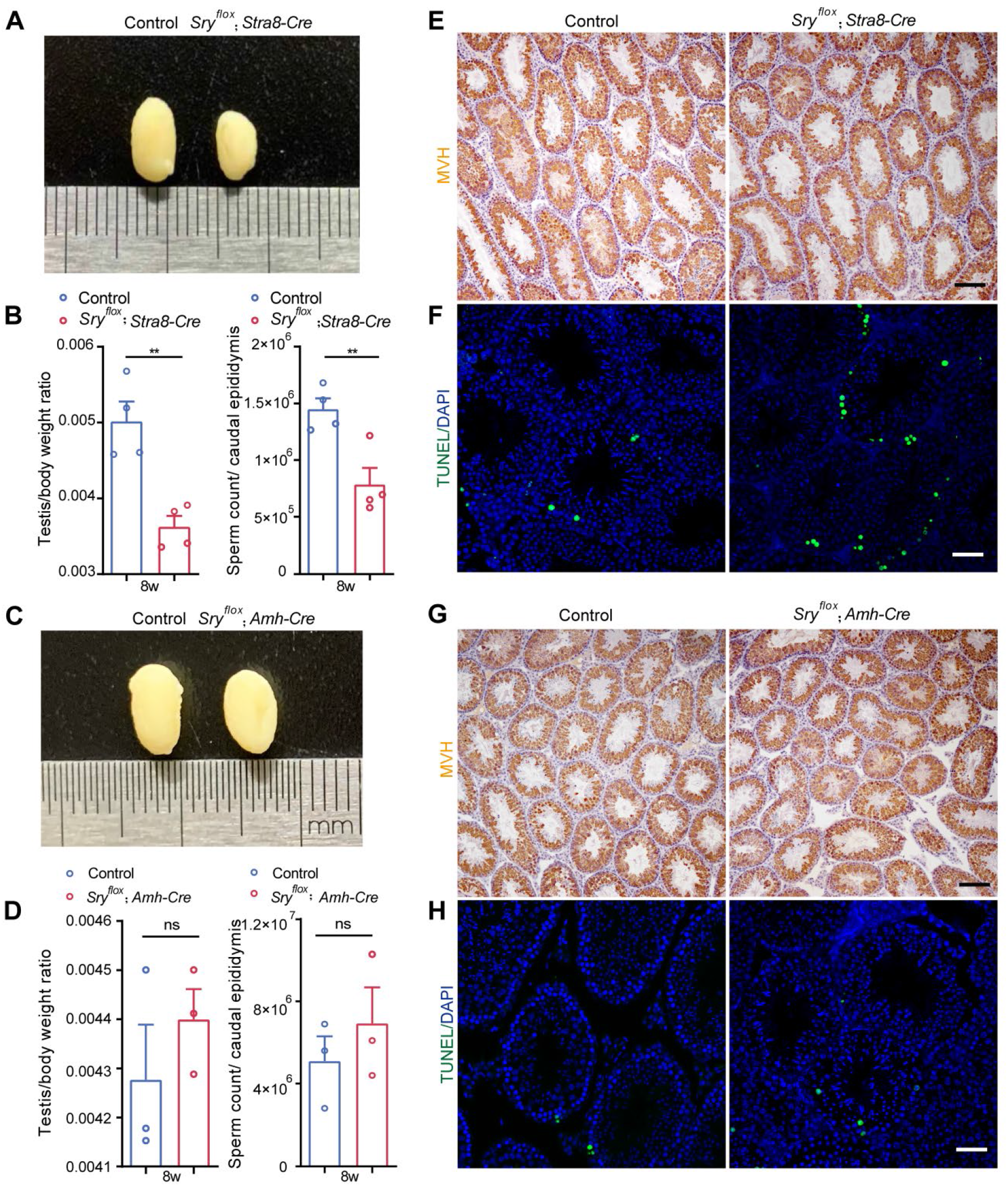
Conditional knockout of *Sry*. (A and C) Representative image of 8 weeks testis from *Sry^flox^*; *Stra8-Cre* or *Sry^flox^*; *Amh-Cre* (right) compared with control mice (left) from the same litter. (B and D) Testis/body ratio and sperm count of *Sry^flox^*; *Stra8-Cre* or *Sry^flox^*; *Amh-Cre* compared with control mice from same litter. (**p<0.01; unpaired, two tailed t test, n=4). (E and G) Germ cells were labeled with antibody against MVH. Loss of epithelium within the seminiferous tubules in *Sry^flox^*; *Stra8-Cre* or *Sry^flox^*; *Amh-Cre* (left) compared with control mice (right). Scale bars indicate 100 μm. (F and H) Tunel assay in *Sry^flox^*; *Stra8-Cre* or *Sry^flox^*; *Amh-Cre* (left) and control mice (right) testis seminiferous tubules. Scale bars indicate 50 μm. The data are presented as the mean ± s.e.m. Source data is available as a Source Data file.

### Loss of *circSry* caused reduction of primary spermatocytes during spermatogenesis

To further characterize the defects of spermatogenesis in *circSry* KO mice, we examined the expression of meiosis-associated genes by immunofluorescence (IF). PLZF positive and STRA8 positive germ cells were localized at the periphery of seminiferous tubules, and no difference was detected in the number of either PLZF positive or STRA8 positive germ cells between control and *circSry* KO testes (Figure 4 figure supplement 1, A and B). Notably, the number of SYCP3-positive germ cells in the seminiferous tubules of *circSry* KO mice was reduced compared with that in control mice (Figure 4, A and B). To assess the progression of spermatogenesis, flow cytometry was used to analyze the proportion of cells with different ploidy levels in 2 month-old testes. The proportion of 4n cells was reduced in *circSry* KO mice, while no difference was noted in the proportion of diploid cells between control and the *circSry* KO mice (Figure 4C). Moreover, TUNEL and SYCP3 double positive germ cells were observed in the *circSry* KO seminiferous tubules, but not in control mice (Figure 4D, a to h). These results suggest that lack of *circSry* leads to defects in meiosis.

**Figure 4.**
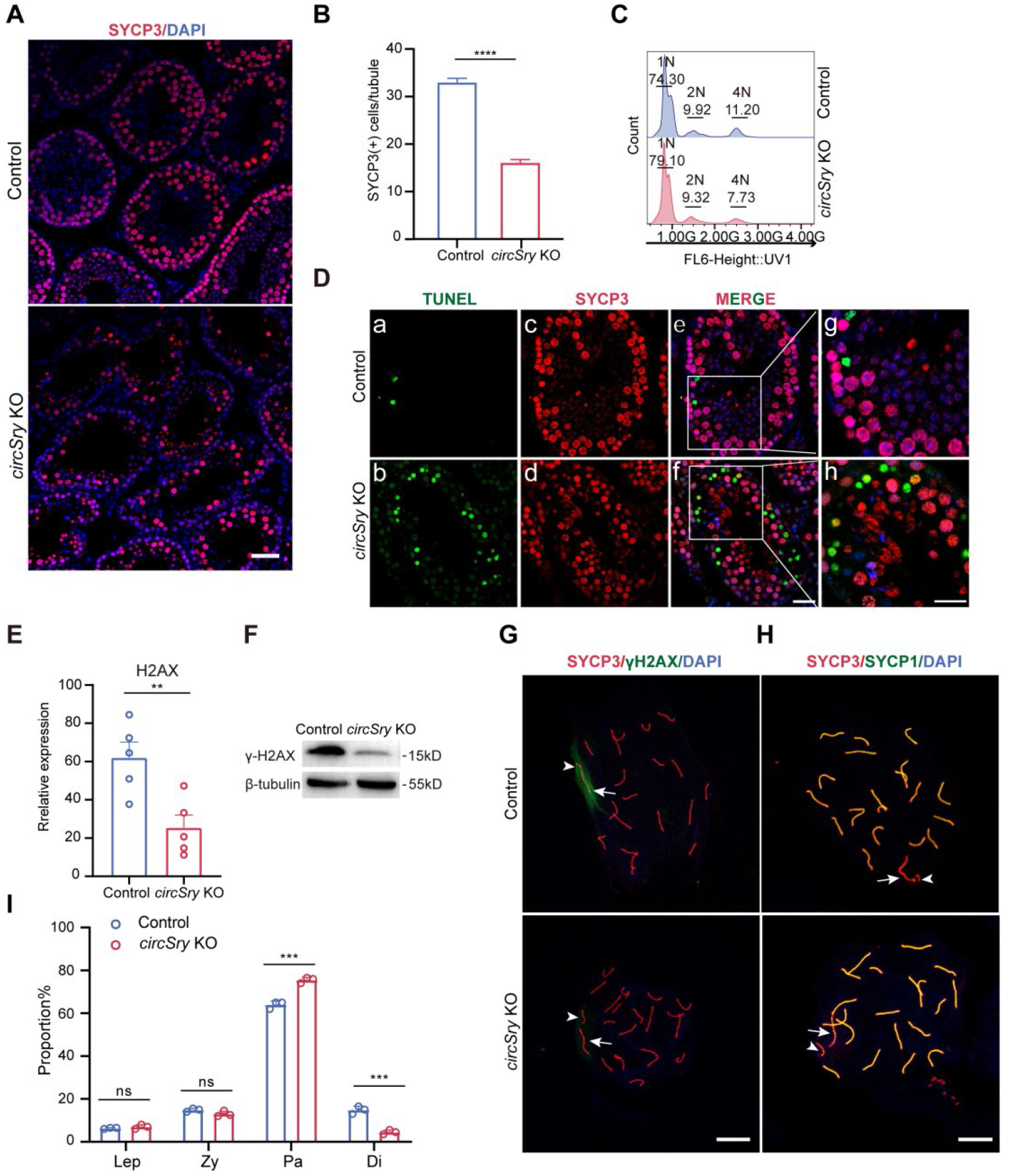
Loss of *circSry* leads to decreased number and meiotic arrest in primary spermatocytes. (A) Immunofluorescence staining of SYCP3 in seminiferous tubules of *circSry* KO (down) and control mice (up). Scale bars indicate 50μm. (B) Quantification of SYCP3 positive cells within seminiferous tubules from *circSry* KO compared with control mice (****p< 0.0001; unpaired, two tailed t test, n=70). (C) Flow cytometry analysis of proportion of 4n cells, 1n cells and 2n cells in 8-weeks mice testes between control and *circSry* KO mice(n=3). (D) (a to h)Representative image of Immunofluorescence co-staining of Tunel signal (green) and SYCP3 (red) in 9-week old control or *circSry* KO mice testes.(a to f Scale bars indicate 50 μm; g, h Scale bars indicate 25 μm).(E) Expression of H2AX mRNA in control and *circSry* KO mice spermatocytes (**p<0.01; unpaired, two tailed t test, n=5) (F) Representative image of western blot results of γH2AX in control and *circSry* KO mice spermatocytes (n=3). (G and H) Immunofluorescence staining of SYCP3 (red) and γH2AX (green) or SYCP1 (green) in control and *circSry* KO spermatocytes at early pachytene stage. Arrowhead indicates the Y chromosome; arrow indicates the X chromosome. Scale bars indicate 100μm. (I)The proportion of leptotene, zygotene, pachytene and diplotene spermatocytes from control or *circSry* KO mice testes. The data are presented as the mean ± s.e.m.

Since it has been reported that loss of histone γH2AX resulted in aberrant synapsis of sex chromosomes during pachynema, which led to meiotic arrest and apoptosis (25, 26), we conducted chromosome spread experiment of spermatocytes with immunostaining of SYCP3 andγH2AX. As shown in Figure 4G, γH2AX expression were decreased in 90% of pachytene stage of *circSry* KO nucleus (n=20) (Figure 4G). To further assess the X-Y synapsis during early pachynema, immunofluorescence staining of SYCP1 and SYCP3 was performed with chromosome spread. In 15.3% of *circSry* KO nucleus (n=300), we observed that X and Y chromosomes paired but not synapsed (Figure 4H). These abnormalities observed in *circSry* KO germ cells were similar with that in *H2ax*^-/-^ germ cells, which led to genomic instability and cell apoptosis (26). Indeed, we observed a significant increase of number of germ cells in pachytene stage and decrease in number of diplotene stage cells (Figure 4I). In contrast, the number of germ cells at leptotene and zygotene stages appeared normal (Figure 4I and Figure 4 figure supplement 3), suggesting that *circSry* functions specifically in pachytene stage.

Failure of XY body formation usually abolishes Meiotic sex chromosome inactivation (MSCI) and increases the expression of X-linked and Y-linked genes in spermatocyte (25). To evaluate whether the deficiency of *circSry* impacted MSCI, we calculated the average value of gene expression from individual chromosomes based on our RNA-seq analysis in *circSry* KO versus control. The average value of all autosomal genes expression was 1.04 (Figure 4 figure supplement 3D), showed that the genes expression level of all autosome chromosomes from *circSry* KO spermatocytes was not different from control. However, the average values of X and Y-linked genes were 1.93 and 2.15, respectively (Figure 4 figure supplement 3D). These data indicated that *circSry* deficiency resulted in the failure of inactivating some of the sex chromosome-linked genes in *circSry* KO spermatocytes. In particular, X-linked genes *Rbbp7*, *Pgk1* (27, 28), and Y-linked gene *Eif2s3y*, known to be subjected to MSCI (25, 29), were up regulated in *circSry* KO spermatocytes, whereas the expression of autosome genes *Xpo6* and *Dhrs1* were not affected (Figure 4 figure supplement 3E). Taken together, we conclude that the loss of *circSry* impairs MSCI.

### *CircSry* regulates γH2AX expression via sponging miR138-5p

Since cytoplasmic circRNAs could act as miRNA sponge to regulate gene expression indirectly, we predicted potential miRNAs that interacted with *circSry* using web tool miRDB (30, 31). 33 miRNAs were predicted to potentially interact with *circSry*, 6 of which had more than 7 binding sites on *circSry*, including well characterized miR-138-5p (Figure 5A and Table S1) (13). To validate these miRNAs using luciferase assay, we constructed reporter plasmid by inserting *circSry* sequence into the 3’UTR of luciferase coding sequence. Compared with scramble miRNA, miR-138-5p, miR-323-5p, and miR-683 reduced the luciferase signals to the greatest extent (Figure 5B). Furthermore, among these miRNAs, miR-138-5p was the most abundantly expressed in mouse male germ cells based on published miRNA sequencing data (32, 33). Therefore, we hypothesized that *circSry* regulated histone H2AX by sequestering miR-138-5p.

**Figure 5.**
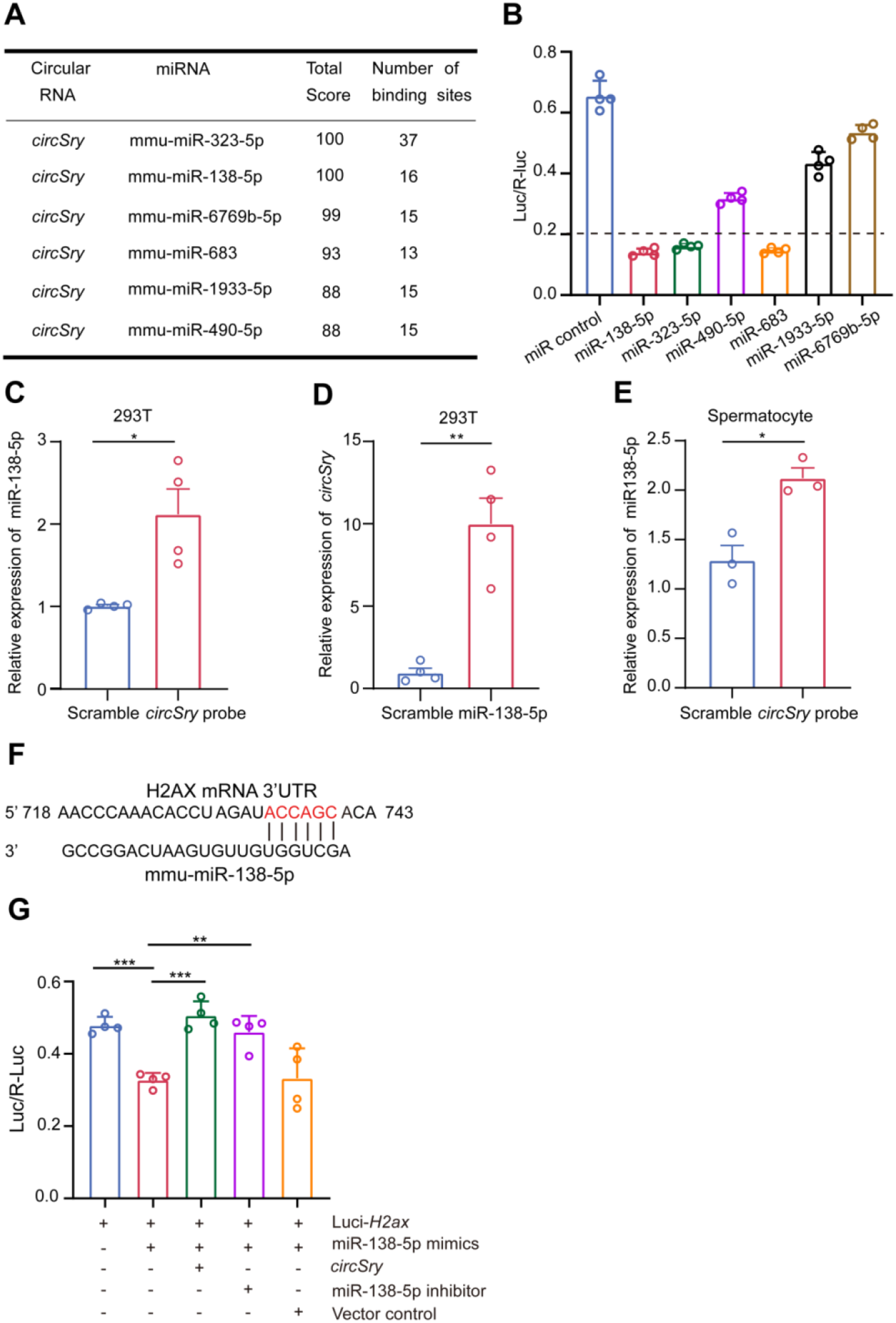
*CircSry* enhances γH2AX expression by sponging miR138-5p in spermatocytes. (A) Table of 6 miRNAs that have more than 7 potential binding sites for *circSry*. (B) Luciferase reporter assay showing diverse binding capacities of 6 miRNAs on Luc-*Sry* (n=4) (C) RNA-RNA pull down enrichment of miR-138-5p using *circSry* biotinylated probe or scramble probe in 293T cell line. (*p<0.05;unpaired, two tailed t test, n=4).(D) RNA-RNA pull down enrichment of *circSry* using biotin-coupled miR-138-5p in 293T cell line. (**p<0.01; unpaired, two tailed t test, n=4). (E) Expression level of miR-138-5p pulled down by *circSry* biotinylated probe or scramble probe in spermatocytes (*p < 0.05, unpaired, two tailed t test n=4). (F) A putative binding site between miR-138-5p and H2AX mRNA 3’UTR. (G) Luciferase reporter assay of *circSry* on sponging miR-138-5p. MiR-138-5p inhibitor and empty vector were used as positive and negative control, separately (**p<0.01, ***p<0.001; unpaired, two tailed t test; n=4). The data are presented as the mean ± s.e.m.

To test whether *circSry* binds AGO2-miR-138-5p complex, we performed AGO2 immunoprecipitation on isolated spermatocytes. *CircSry* was specifically enriched in AGO2 immunoprecipitates (Figure 5 figure supplement 1A), indicating that *circSry* interacts with miRNAs. Next, we performed RNA-RNA pull down experiment to examine the interaction between miR-138-5p and *circSry* in 293T cell line that highly expressed *circSry* (Figure 5 figure supplement 1B). This cell line was established by transducing a lentiviral vector harboring the *circSry* expressing cassette. The splicing site of *circSry* was also confirmed by Sanger sequencing (Figure 5 figure supplement 1C). Pull down experiment showed that *circSry* biotinylated probe significantly enriched the miR138-5p compared to the scramble probe (Figure 5C), and biotin-coupled miR-138-5p captured more *circSry* than the biotin control miRNA (Figure 5D). Furthermore, RNA-RNA pull down in spermatocytes showed that *circSry* probe captured more miR-138-5p than scramble probe as well (Figure 5E). Taken together, these results suggest that *circSry* acts as a sponge for miR-138-5p.

It has been reported that miR-138-5p directly down regulates H2AX expression through binding to 3’UTR region of H2AX mRNA (Figure 5F), inducing chromosomal instability during DNA damage repair (34). To test our hypothesis that *circSry* regulates H2AX expression via sponging miR-138-5p during spermatogenesis, we conducted luciferase reporter experiments. Transfection of luciferase reporter containing H2AX sequence together with miR-138-5p mimics showed significantly reduced luciferase signal, while co-transfection of *circSry* rescued the decrease of luciferase signal (Figure 5G).

Taken together, these results show that *circSry* regulates the expression of H2AX mRNA at post-transcriptional level by sponging miR-138-5p and contributes to MSCI during spermatogenesis.

## Discussion

Recent studies have identified thousands of circular RNAs and many of them have important biological functions in various tissues and organs. Here we show that a circular transcript exhibits regulatory function in male reproductive system. Even more interesting, this circular RNA originates from the sex-determining gene *Sry*, which initiates the male germ cell development in the first place. In this study, we show that loss of *circSry* led to defective spermatogenesis, specifically causing primary spermatocytes apoptosis. Deletion of *circSry* decreased the expression of H2AX and displayed aberrant XY synapsis at pachytene stage, followed by MSCI abolishment. Given that miR-138-5p directly targets H2AX mRNA (34), we performed RNA-RNA pull down experiment and luciferase reporter assay, demonstrating that *circSry* enhanced H2AX expression by sequestering miR-138-5p.

Mouse *Sry* gene is capable of producing linear RNAs at embryonic stage as well as circular transcripts in adult testis. It was proposed that *circSry* is generated from a linear RNA precursor containing long palindromic repeats, which is transcribed from a distal promoter (35). How does the transcription of *Sry* differentially regulated in embryonic genital ridge and adult male germs cells is poorly understood. This mechanism of alternative promoter choice and upstream regulation is an interesting direction of future study.

In addition to *Mus musculus*, transcripts of *Sry* were also detected in the testes of *Mus musculus domesticus* and *Mus spretus*. Considering that splicing donor site was also conserved in *Rattus norvegicus* (21), we speculated that alternative splicing occurred in rat as well. Do these transcripts form circular RNA, and do they regulate spermatogenesis need to be further explored. *CircSry* was also expressed in the mouse brain during embryonic stage and its expression was diminished after birth (36). Whether *circSry* plays a role in mouse brain needs further investigation.

Human *SRY* is also expressed in adult testis. Different from the circular transcript in adult mouse testis, however, human *SRY* transcript is liner and polyadenylated (37), and the presence of circular *SRY* has not been reported. The function of human *SRY* transcript in adult testes is very curious and the elucidation of which may help us to better understand male development and spermatogenesis in human. While human *SRY* transcript in testes might regulate germ cell development using a different mechanism, the sex determining gene also plays a role in adult testes could be a common theme in multiple mammalian species.

Taken together, our study complements *Sry*’s role in germ cell development, revealing its significance in male germ cell development. *CircSry* safeguards the inactivation of sex chromosomes during pachytene stage, the key process to complete meiosis and produce sperms. Upon fertilization, the sperm carrying Y chromosome contributes Y to the embryo which will develop into a male and grow testicles. The cycle of *Sry* expression and duel forms and functions suggests a mechanism to ensure the preservation of vulnerable Y chromosome in evolution.

## Materials and Methods

### Animals

The mice used in this research were C57BL/6 background. All animals were maintained at 24°C and 50–60%humidity under a 12:12 h light/dark cycles and with ad libitum access to food and water. *Sry^flox^* mice were generated in Jackson Laboratory.*Stra8-Cre* and *Amh-Cre* mice were provided by Prof. Gao of Institute of Zoology, University of Chinese Academy of Sciences. All mice studies were carried out in accordance with the principles approved by the Institutional Animal Care and Use Committee at the Institute of Zoology, Chinese Academy of Sciences.

### Establishment of mutant mice by CRISPR/Cas9-based genome editing

*CircSry* knockout mice were established by microinjection of Cas9 mRNA and single-guide RNAs (sgRNAs) into zygotes (38). All RNAs prepared for microinjection were in vitro transcribed. Briefly, Cas9 mRNAs and sgRNAs were mixed properly and injected into zygotes. The injected embryos were transferred to pseudopregnant females. To genotype mutant mouse lines, genomic sequence was amplified by PCR, followed by Sanger sequencing. All oligonucleotides were listed in the Table S2.

### Establishment of *Sry^flox^* mice by CRISPR/Cas9 system

We used single-stranded oligodeoxynucleotides (ssODN) to establish knock-in mouse mutants (38). Establishment of *Sry^flox^* KI mice went through two rounds of microinjections. The first microinjection contained Cas9 mRNA and ssODN (V5-loxP) targeted the 3’ end of *Sry* exon 1(V5-TGA-loxP). The injected embryos were transferred to pseudopregnant females. The correct male mouse line was determined by Sanger sequencing of genomic sequence. Sperms from *Sry*-V5-loxP male mouse line were collected for single-sperm microinjection of wild-type oocytes. The second microinjection contained Cas9 mRNA and ssODN (5’-loxP) targeted the 5’ upstream site of *Sry* exon1 start codon within the *Sry*-V5-loxP zygotes. The correct mouse line was again confirmed by Sanger sequencing of genomic sequence. All oligonucleotides and primers sequences were listed in the Table S2.

### Genotyping of mice and sexing

All mice were genotyped with the tail DNA. Chromosomal sex was detected through amplifying the Y-linked gene *Uty*. Phenotypic sex was determined by examination of external genitalia, and the presence /absence of mammary glands. Primers were listed in Table S2.

### RNA preparation and real-time PCR

Nuclear and cytoplasmic RNAs were extracted by using Norgen’s Cytoplasmic & Nuclear RNA Purification Kit (Norgen Biotek Corp.). RNAs were extracted using Trizol reagent (Life Technologies). cDNAs were synthesized using the Hifair III 1^st^ strand cDNA synthesis reagent (Yeasen company, China) with 500ng of total RNA. cDNAs were amplified with Hieff qPCR SYBR Green Master mix (Yeasen company, China) and quantified with Roche LightCycle 480 system. For RNase R treatment, experiment was performed by incubation of 3 μg of RNA with 6U μg^-1^ of RNase R (Epicenter) for 25 minutes at 37°C. The expression levels of each gene were presented relative to GAPDH or U6 expression. The primers sequences were listed in Table S2.

### RNA pull-down

Biotinylated *circSry* probe and 5’bio-miRNA mimic were synthesized by RiboBio (Guangzhou, China). Testes were ground and incubated in lysis buffer [50 mM Tris-HCl, pH 7.4, 150 mM NaCl, 2 mM MgCl_2_, 1% NP40, RNase Inhibitor (Beyotime biotechnology)] on ice for 1 hour. The lysates were then incubated with the biotinylated probes at RT for 4 hours; followed by adding streptavidin C1 magnetic beads (Invitrogen) were to binding reaction, and continued to incubate at 4°C, overnight. On second day, the beads were washed briefly with wash buffer [0.1% SDS, 1% Triton X-100, 2 mM EDTA, 20 mM Tris-HCl, and 500 mM NaCl] five times. The bound RNA in the pull-down was further extracted for purification and RT-qPCR. The sequence of *circSry* probe was shown in Table S3.

### RNA-binding protein immunoprecipitation

RIP experiments were performed with primary spermatocytes which separated from adult mice testes and homogenized into a single-cell suspension in ice-cold PBS. After centrifugation, the pellet was resuspended in RIP lysis buffer. Magnetic beads were incubated with 5 μg antibody against AGO2 (Proteintech Inc), or IgG at RT. The tissue lysates were then incubated with the bead-antibody complexes overnight at 4°C. RNAs were extracted by Trizol reagent and reverse-transcribed after proteinase K treatment.

### RNA-seq and data analysis

RNA sequencing reads were aligned to mouse reference sequence GRCm39 using STAR (2.7.0f). Read counts and TPM (Transcripts Per Kilobase Million) were counted using RSEM (1.3.2). Differential expression genes (DEG) (FDR < 0.05, log2 (fold change) (log2 FC) ≥ 1 or ≤ −1) were calculated by edgeR (3.28.1) package. The DEG genes were carried out Gene Ontology Pathway analysis and Kyoto Encyclopedia of Genes and Genomes analysis by clusterProfiler (3.14.3) and org.Hs.eg.db (3.10.0) packages. Heatmap (1.0.12) was used for the heatmap visualization; colors represent the Z-score derived from the log2 transformed FPKM data. Principal Component Analysis (PCA) was performed by R packages, including tidyr (1.1.2), dplyr (1.0.2) and ggplot2 (3.3.2).

### Construction of *circSry* expression vector

The empty vector was purchased from Geneseed Bio (Guangzhou,China), termed pLC5.This plasmid was linearized with XhoI-BamHI. Complete *circSry* sequence was amplified with XhoI-BamHI clone site and inserted into pLC5.The final plasmid was named pLC5-*circSry*. After sequencing, pLC5-*circSry* was transfected into 293T cell line to testify the expression of *circSry*.

### Dual-luciferase reporter assay

The full-length sequence of *circSry* or the 3’UTR of H2AX was inserted into the 3’ UTR of pMIR-Report Luciferase vector (gifted by Yu Wang lab, State Key Laboratory of Stem Cell and Reproductive Biology, Institute of Zoology, Chinese Academy of Sciences, Beijing). Co-transfection of 500 ng pMIR-Report luciferase vector, miRNA mimic (RiboBio) and pLC5-*circSry* were conducted using lipofectamine 2000 (Invitrogen). After 48 hours, the luciferase activities were measured using a dual-luciferase reporter assay kit (Promega). The results were normalized to the ratio between Firefly signal and Renilla signal.

### Western blotting

Protein obtained from spermatocytes were separated by gel electrophoresis SDS-PAGE and transferred to a PVDF membrane. The PVDF membrane was incubated with primary antibodies against γH2AX (Millipore; 05-636, 1:200), β tubulin (Yeasen company, China; 1:1000) 4°C overnight and horseradish peroxidase-labeled secondary antibody for 1 hour at 37°C subsequently. Images were captured using ECL Western Blotting Substrate (Thermo Scientific Pierce).

### Tunel assays

The TUNEL assay was performed by TUNEL BrightRed Apoptosis Detection Kit (Vazyme). Briefly; sections were permeabilized by protein K and labeled with rTdT reaction mix for 1 hour at 37°C. Reaction was stopped by 1× PBS. After washing in PBS, the sections were incubated in 2 μg/ml of DAPI (Molecular Probes, D1306) for 10 minutes. Sections were mounted on slices with Dako Fluorescence Mounting Medium (Dako Canada, ON, Canada). Images were obtained using a laser scanning confocal microscope LSM780 (Carl Zeiss).

### Flow cytometry

The cell suspensions obtained from testes were digested by collagenase IV and Typsin for 5 minutes (39). To analyse DNA ploidy, the cells were incubated with the Hoechst for 15 minutes and filtered before being subjected to flow cytometry. The results were analysed using a FACS-Calibur system (BD Biosciences).

### Sperm count and fertility

The epididymis tail of the mouse was taken out, cut into pieces with ophthalmic scissors, placed in a 37°C water bath for 15 minutes, and counted via microscope. 2-3-month-old *circSry* and control C57BL/6J male mice were housed with control C57BL/6J males (2–3-month-old), which were proved having normal fecundity. Copulatory plugs were monitored daily, and plugged females with visibly growing abdomen were moved to separate cages for monitoring pregnancy. The mating process lasted for 4 months. The numbers of pups (both alive and dead) were counted on the first day of life.

### Tissue collection and histological analysis

Testes from control and *circSry* KO mouse were dissected immediately after euthanasia, then immediately fixed in 4% paraformaldehyde (PFA) for 24 hours, after storing in 70% ethanol and embedding in paraffin, 5 μm-thick sections were prepared using a rotary microtome (Leica) and mounted on glass slides. Sections were stained with MVH (Abcam) for histological analysis.

### Immunofluorescence analysis

After deparaffinization and antigen retrieval, 5% bovine serum was used to block sections at room temperature (RT) for 1 hour, and specific primary antibody was used to incubate with sections at RT or overnight at 4°C. After washing the sections for three times, the slides were incubated with the corresponding secondary antibody, fluorescent dye-conjugated-FITC or TRITC (1:150, Jackson) for 1 hour at RT (Avoiding the light). DAPI was used to stain the nucleus. All images were captured with a confocal microscopy (Leica TCS SP8). All the antibodies were listed in the methods and materials.

### Preparation of synaptonemal complex

Seminiferous tubules were collected from dissecting testes and washed with 1× PBS. Hypo extraction buffer (HEB) was used to incubate within seminiferous tubules for 30 minutes, followed by disrupting within 0.1 M sucrose liquid to form a single-cell suspension. The cell suspension was mounted on slides treated with 1% PFA. Slides were air-dried in a humidified box for at least 6 hours. After washing with 0.04% Photo-Flo (Equl, 1464510), the slides were staining for SYCP3. Incubating with SYCP3 antibody was conducted at RT for 30 minutes after antibody dilution buffer (ADB) treatment. After washing three times in 1× Tris buffer, saline (TBS), blocking was conducted with 1× ADB at 4°C overnight. After washing three times with cold 1× TBS buffer, corresponding secondary antibody, and fluorescent dye-conjugated-TRITC were incubated with sections for 3 hours at 37°C. All images were captured with a confocal laser scanning microscope (Leica TCS SP8).

### Data and statistical analysis

All images were processed with Photoshop CS6 (Adobe). All statistics were analyzed using Prism software (GraphPad Software). All experiments were confirmed with at least three independent experiments, and three to five control or mutant testes were used for immunostaining. The quantitative results were presented as the mean ± s.e.m. The significant difference was evaluated with t-test. P-value < 0.05 was considered as significant.

## Data availibility

RNA sequencing-derived data reported in this study have been deposited in NCBI’s Gene Expression Omnibus (GEO) under accession number GSE185184.Reviewers go to GEO with the link: https://www.ncbi.nlm.nih.gov/geo/query/acc.cgi?acc=GSE185184. A private token is provided for reviewers to check the records: sditsmkadvqvfqh.

Source data is provided as Supplementary file2.

## Acknowledgments

We are grateful to Chenrui An for their comments and editing of the manuscript; and all members of Fei Gao laboratory for technical support. This work was supported by National Key Research and Development Program of China (2019YFA0110000, 2018YFE0201102, 2018YFA0107703 and 2016YFA0101402), Strategic Priority Research Program of the Chinese Academy of Sciences (XDA16010503) Strategic Collaborative Research Program of the Ferring Institute of Reproductive Medicine, Ferring Pharmaceuticals, Chinese Academy of Sciences (FIRMD181101),National Natural Science Foundation of China (31722036 and 81773269).

---

**Figure 1 figure supplement 1.**
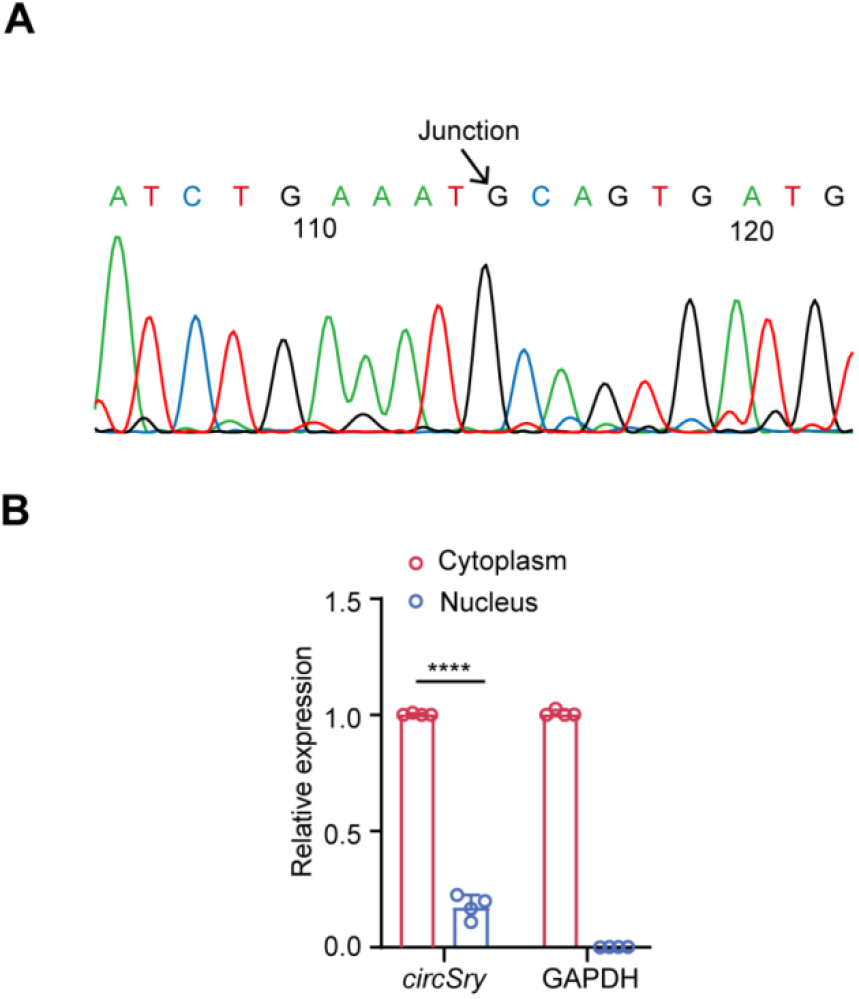
Characterization of *circSry* in mouse testis (related to Figure 1). (A) Splicing junction was confirmed by Sanger sequencing. The arrow indicated the head-to-tail splicing site of *circSry*. (B) qRT-PCR results of cytoplasm-nucleus distribution of *circSry* (****P<0.0001; unpaired two-tailed t test, n=4). The data are presented as the mean ± s.e.m.

**Figure 2 figure supplement 1.**
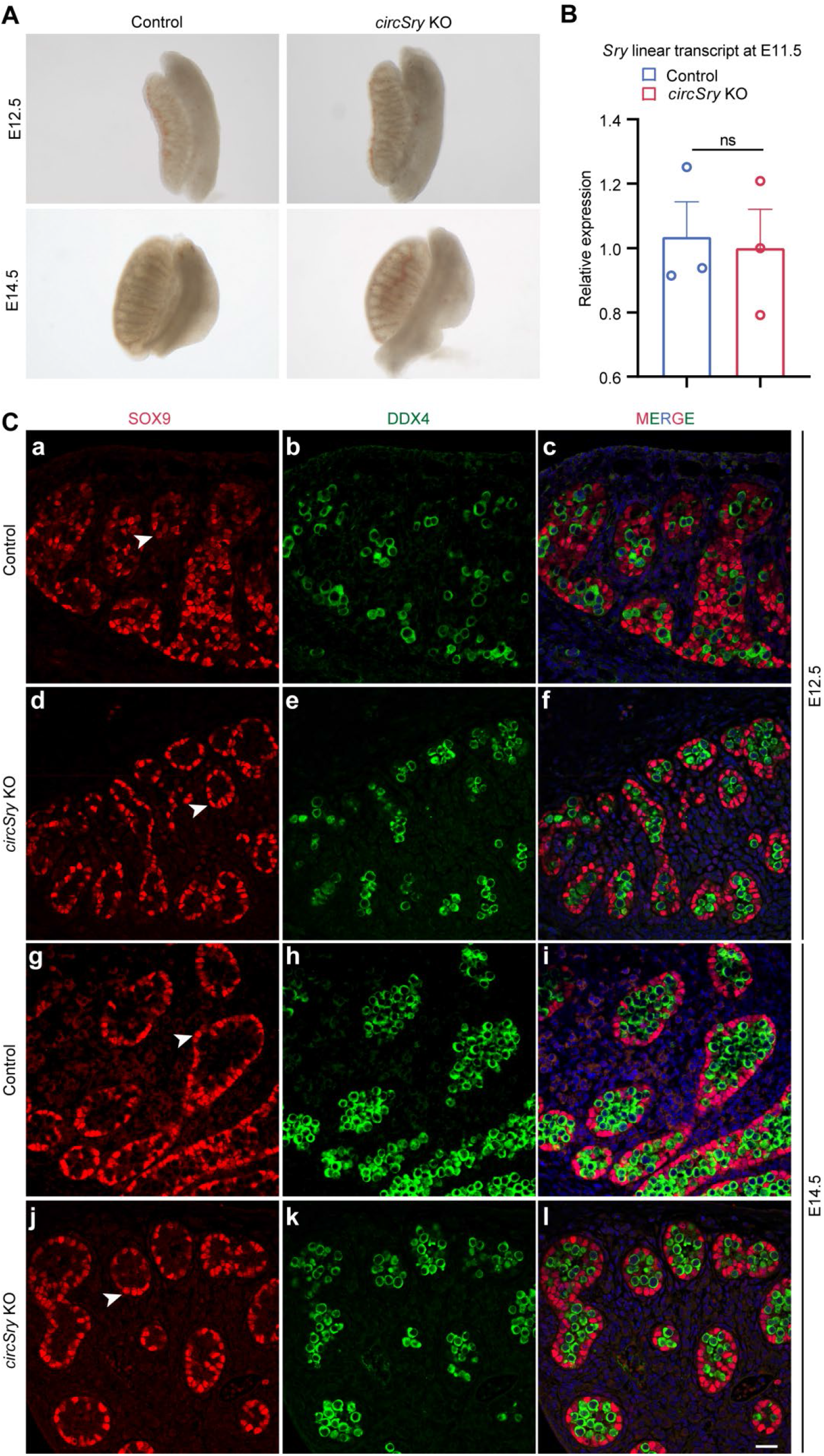
No significant difference was detected in *circSry* KO embryos gonads (related to Figure 2). (A) No significant difference was detected in control and *circSry* KO gonads morphology at E12.5 and E14.5 (B) Quantitative analyses of *Sry* gene in control and *circSry* KO gonads at E11.5. The mRNA expression level of *Sry* in in control and *circSry* KO gonads was not significantly changed at E11.5. The gender of the embryos was confirmed with PCR using *Sry* primers. Data are presented as the mean ± s.e.m; unpaired, two tailed t test, ns, not significant, p > 0.05. (C)SOX9 was expressed in Sertoli cells of both control and *circSry* KO gonads at E12.5 and E14.5. SOX9/DDX4 double-staining experiment was performed with control and *circSry* KO embryos at E12.5 (a–f) and E14.5 (g–l). Germ cells were labeled with DDX4 (green). DAPI (blue) was used to stain the nuclei. The arrowheads point to SOX9-positive Sertoli cells (red). The gender of the embryos was confirmed with PCR using *Sry* primers. Scale bars indicate 50 μm.

**Figure 2 figure supplement 2.**
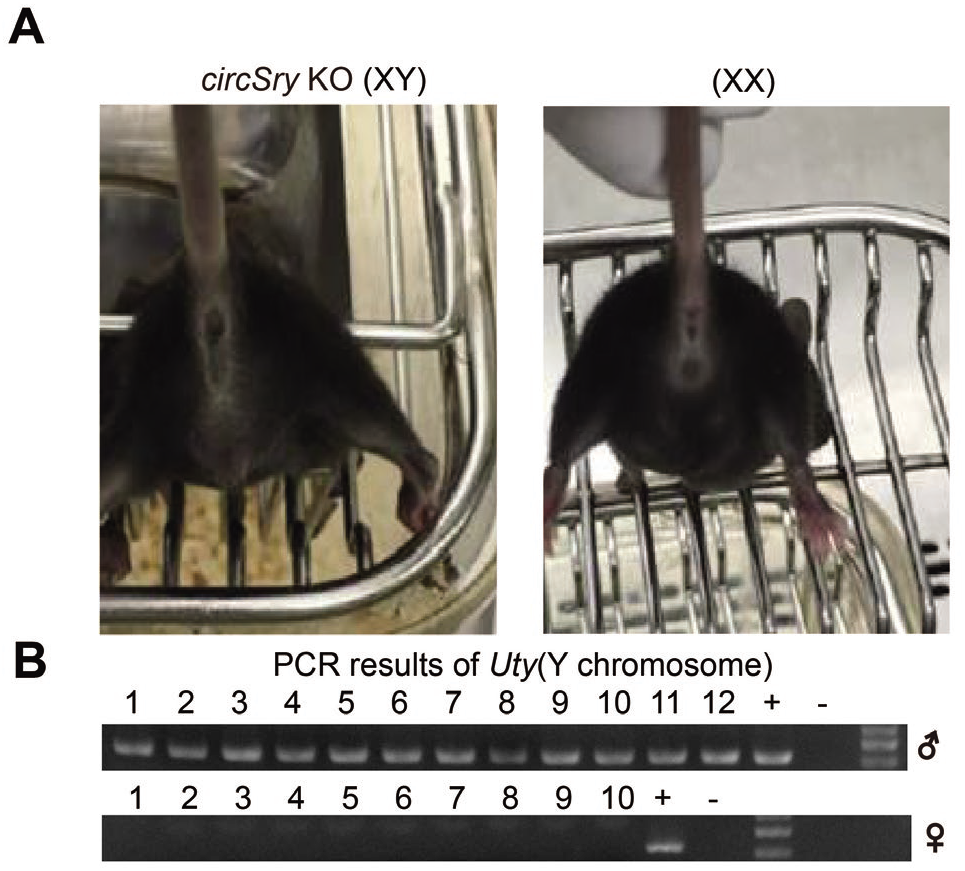
*CircSry* KO mouse develop as male (related to Figure 2). (A) F1 offsprings of *circSry* KO developed as male, and offspring XX mice exhibit normal female external genitalia and mammary glands. (B) PCR results of gender identification of F1 off springs. Y chromosomal gene *Uty* was detected by PCR.

**Figure 2 figure supplement 3.**
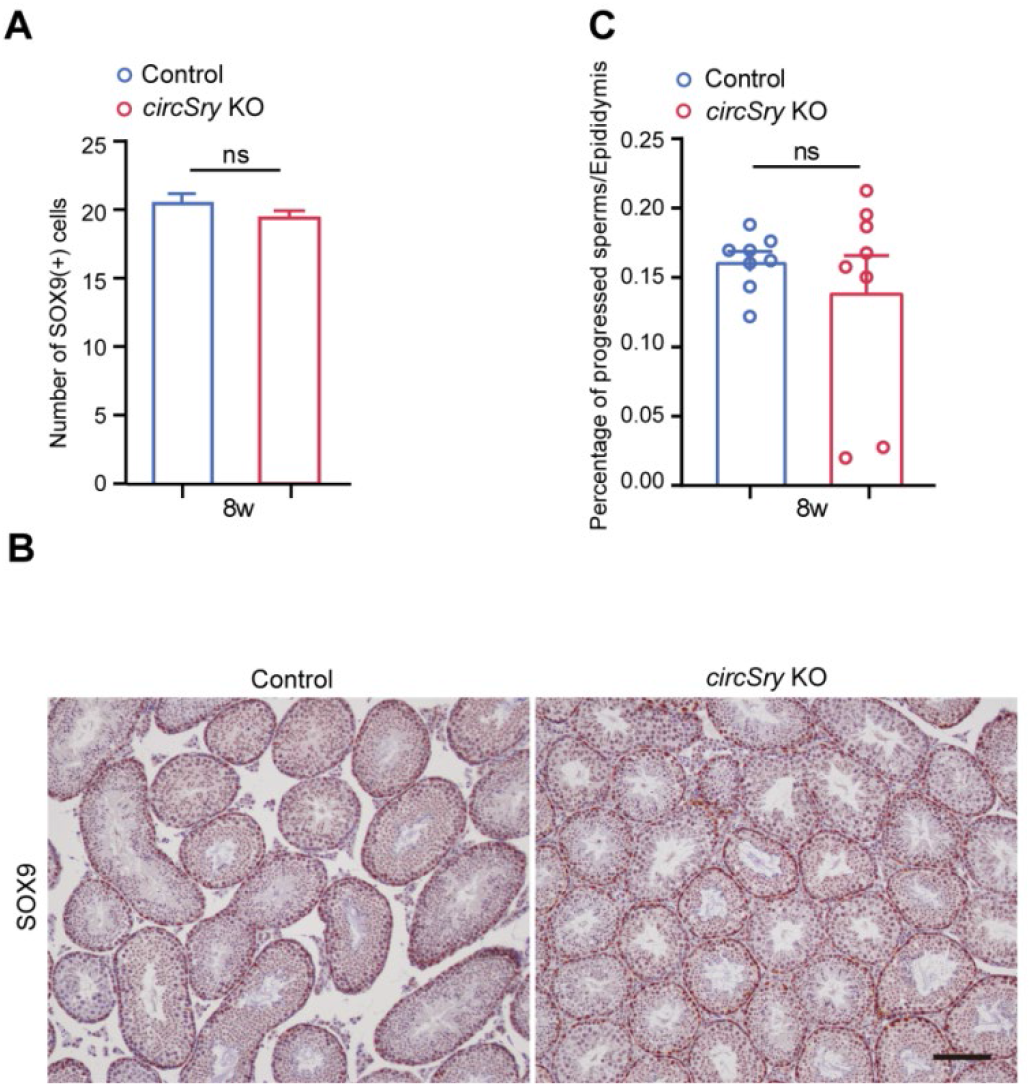
Immunohistochemical staining of SOX9 and sperm motility analysis in control and *circSry* KO mice (related to Figure 2). (A) Quantification of progressed sperm obtained from epididymis between *circSry* KO and control mice. (ns, not significant; unpaired, two tailed t test, n=8). Sperms were collected from 8-week old *circSry* KO or control mice epididymis. (B) Representative image of Immunofluorescence staining of SOX9 in 8-week old control and *circSry* KO mice testes. Scale bars indicate 50 μm. (C) Quantification of SOX9 positive cells within seminiferous tubules from 8-week old control and *circSry* KO mice (ns, not significant; unpaired, two tailed t test, control, n=30; *circSry* KO, n=30). The data are presented as the mean ± s.e.m.

**Figure 3 figure supplement 1.**
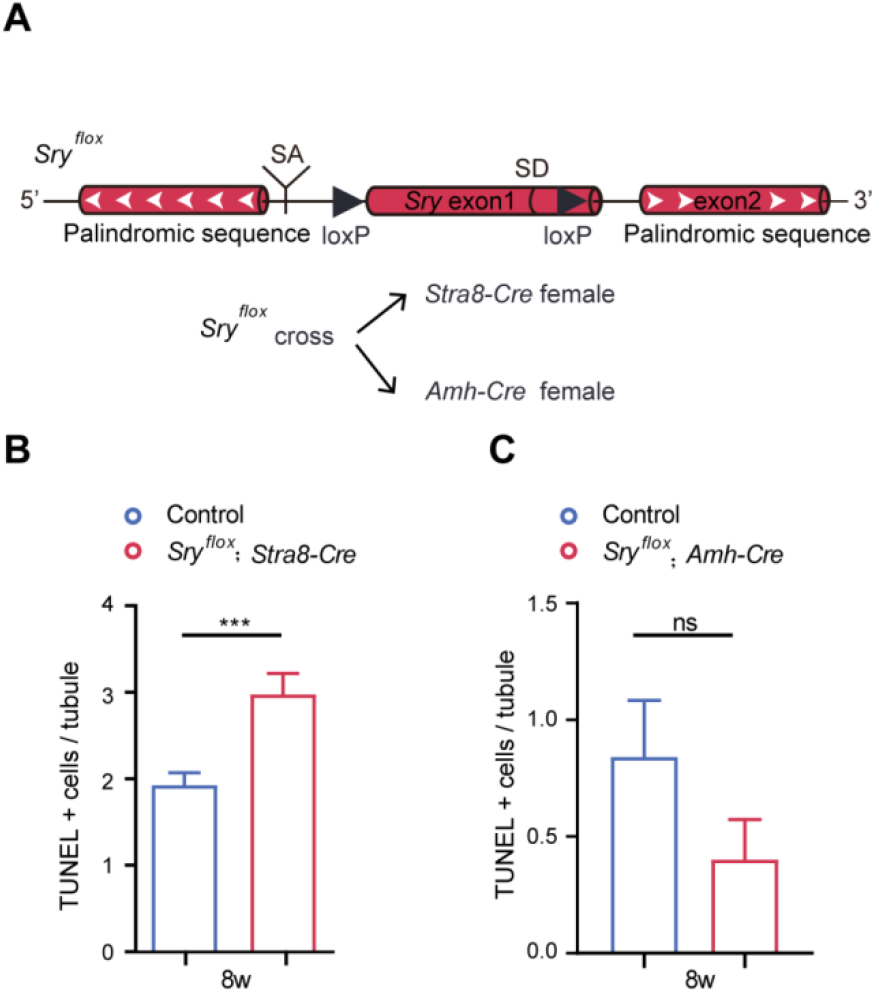
Conditional knockout of *Sry* (related to Figure 3). (A) Schema of inserting loxP sequences into *Sry* locus. *Sry^flox^* was mated with *Stra8-Cre* female or *Amh-Cre* female to generate conditional knockout mice in germ cells or Sertoli cells, respectively. (B and C) Quantification of apoptotic cells in *Sry_flox_*; *Stra8-Cre* or *Sry^flox^*; *Amh-Cre* mice testis seminiferous tubules. The total number of calculated tubules reached 150. (***p<0.001; unpaired, two tailed t test; ns, not significant). The data are presented as the mean ± s.e.m.

**Figure 4 figure supplement 1.**
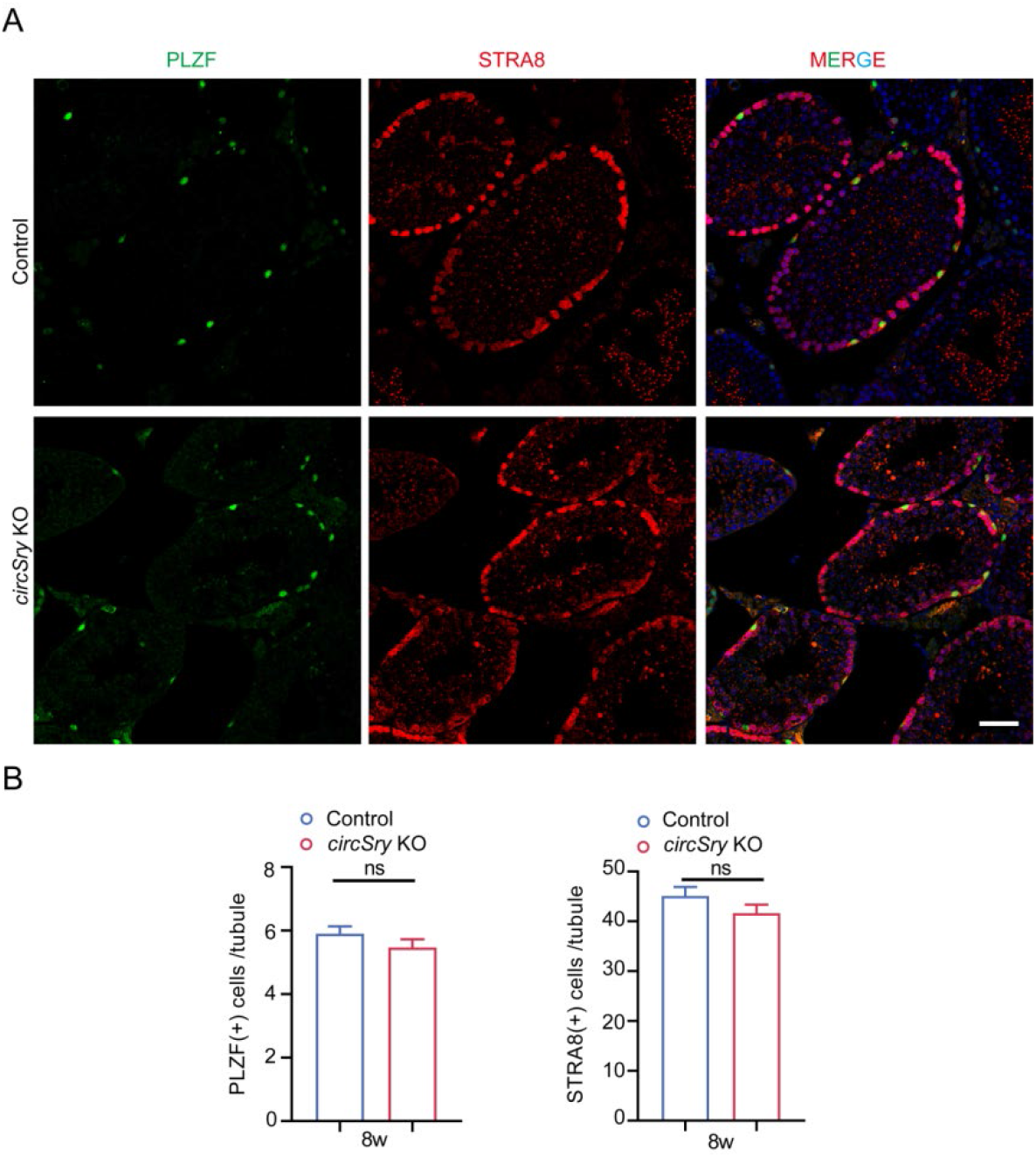
Immunofluorescence co-staining PLZF and STRA8 of control and *circSry* KO testes (related to Figure 4). (A)Representative image of Immunofluorescence staining of PLZF (green) and STRA8 (red) in 8-week old control and *circSry* KO mice testes. Scale bars indicate 50 μm. (B) Quantification of the number of PLZF positive cells (left) and STRA8 positive cells (right) within seminiferous tubules from three independent mice of 8-week old control and *circSry* KO mice (ns, not significant; unpaired, two tailed t test, control, n=30; *circSry* KO, n=30). The data are presented as the mean ± s.e.m.

**Figure 4 figure supplement 2.**
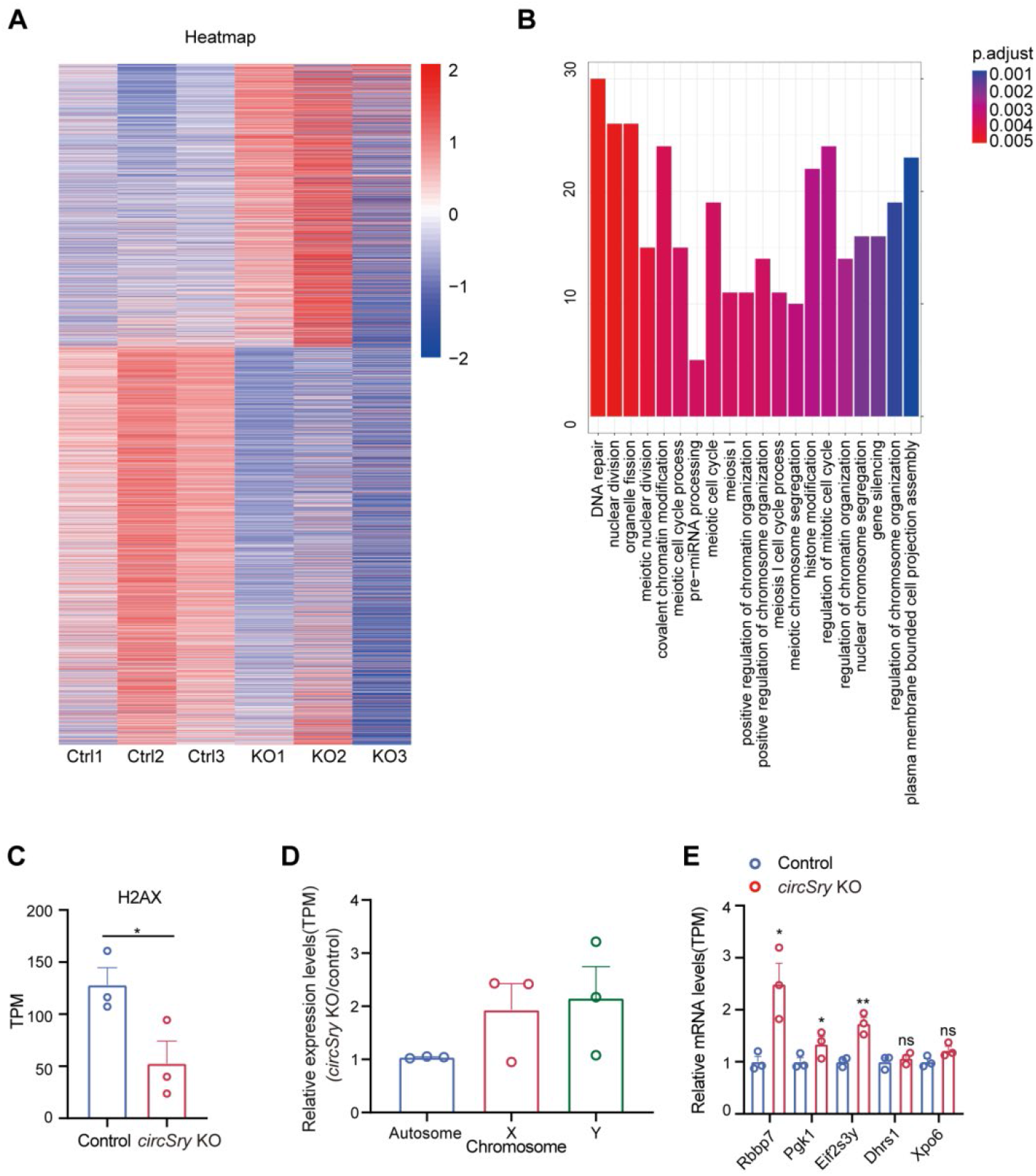
RNA-seq analysis between *circSry* KO and control primary spermatocytes (related to Figure 4). (A) Heat map illustrates RNA-seq differential expression data. Red, positive fold-change (logFC) indicates higher expression; blue, negative logFC. Total RNA was extracted from primary spermatocytes of control or *circSry* KO mice. (B) Enrichment analysis of significant down regulation comparing with control mice, FDR<0.05, logFC<-1. (C) H2AX expression comparing with control and *circSry* KO spermatocytes (*p<0.05, t test, n=3). (D) Relative gene expression levels on chromosomes as determined by RNA-seq (n=3). (E) Relative expression of X-linked genes, *Rbby7*, *Pgk1*; Y-linked genes, *Eif2s3y*; and genes on autosome, *Xpo6*, *Dhrs1*, as determined by RNA-seq analysis on control and *circSry* KO spermatocytes (**p<0.01, *p<0.05, ns, not significant; n=3).The data are presented as the mean ± s.e.m.

**Figure 4 figure supplement 3.**
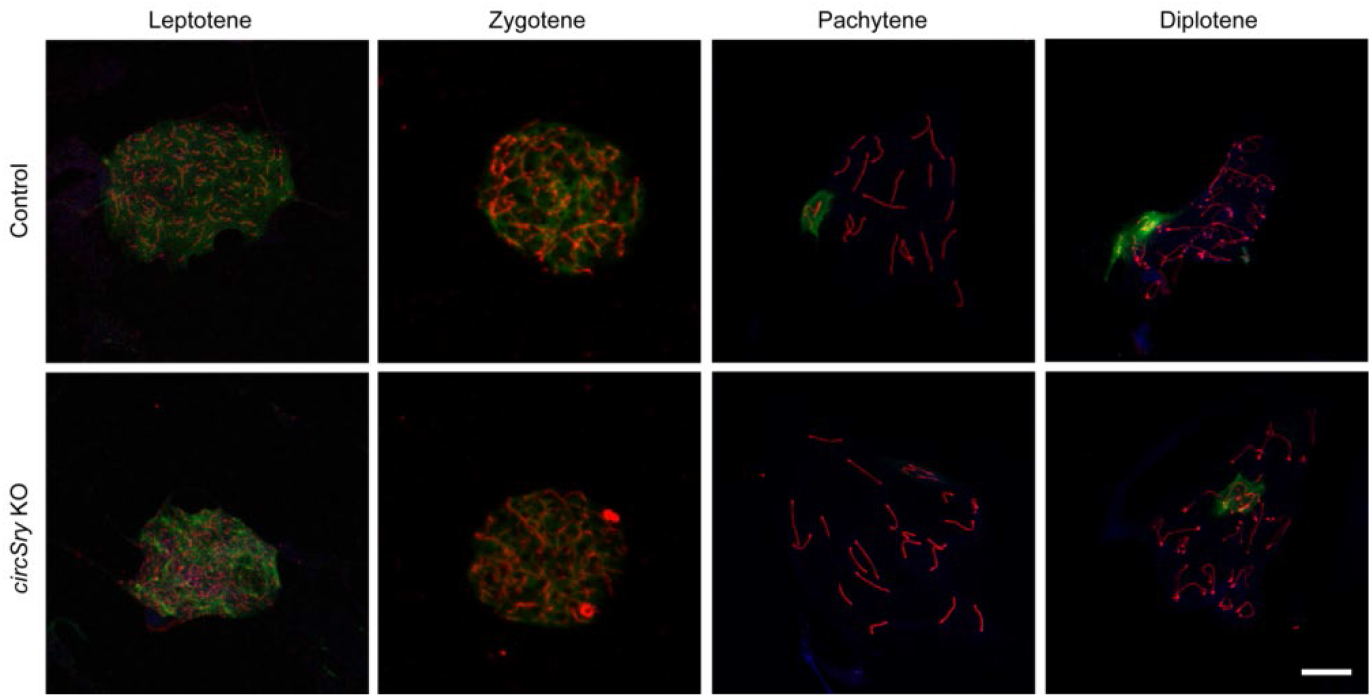
Staining SYCP3 (red) and γH2AX (green) in control and *circSry* KO spermatocytes from leptotene to diplotene stages (related to Figure 4). (Up) Representative image of immunofluorescence staining of SYCP3 (red), γH2AX (green) and DAPI (blue) in 8-week old control or *circSry* KO mice testes (Down). Scale bars indicate 10 μm. (Two tailed t test, ***p<0.001, ns, not significant). Total 400 cells (300 cells of pachytene stage, 100 cells of the other stages) were counted from 3 mice.

**Figure 5 figure supplement 1.**
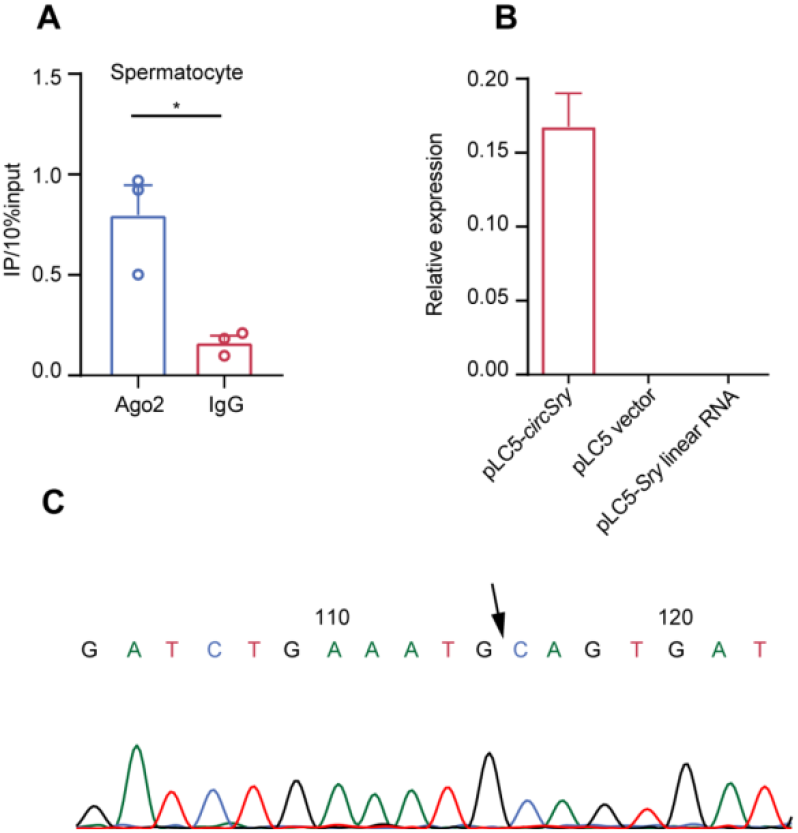
Overexpression of *circSry* in 293T cell line (related to Figure 5). (A) RIP results showing the *circSry* enrichment using AGO2 antibody vs IgG antibody *in vivo* (*p< 0.05; unpaired, two tailed t test, n=3). (B) Relative expression of *circSry* and *Sry* linear RNA were measured with 293T cells line that expressed *circSry*. (C) Sanger sequencing showed junction sequence of *circSry* (Arrow indicates the junction site). The data are presented as the mean ± s.e.m.

**Table S1.**
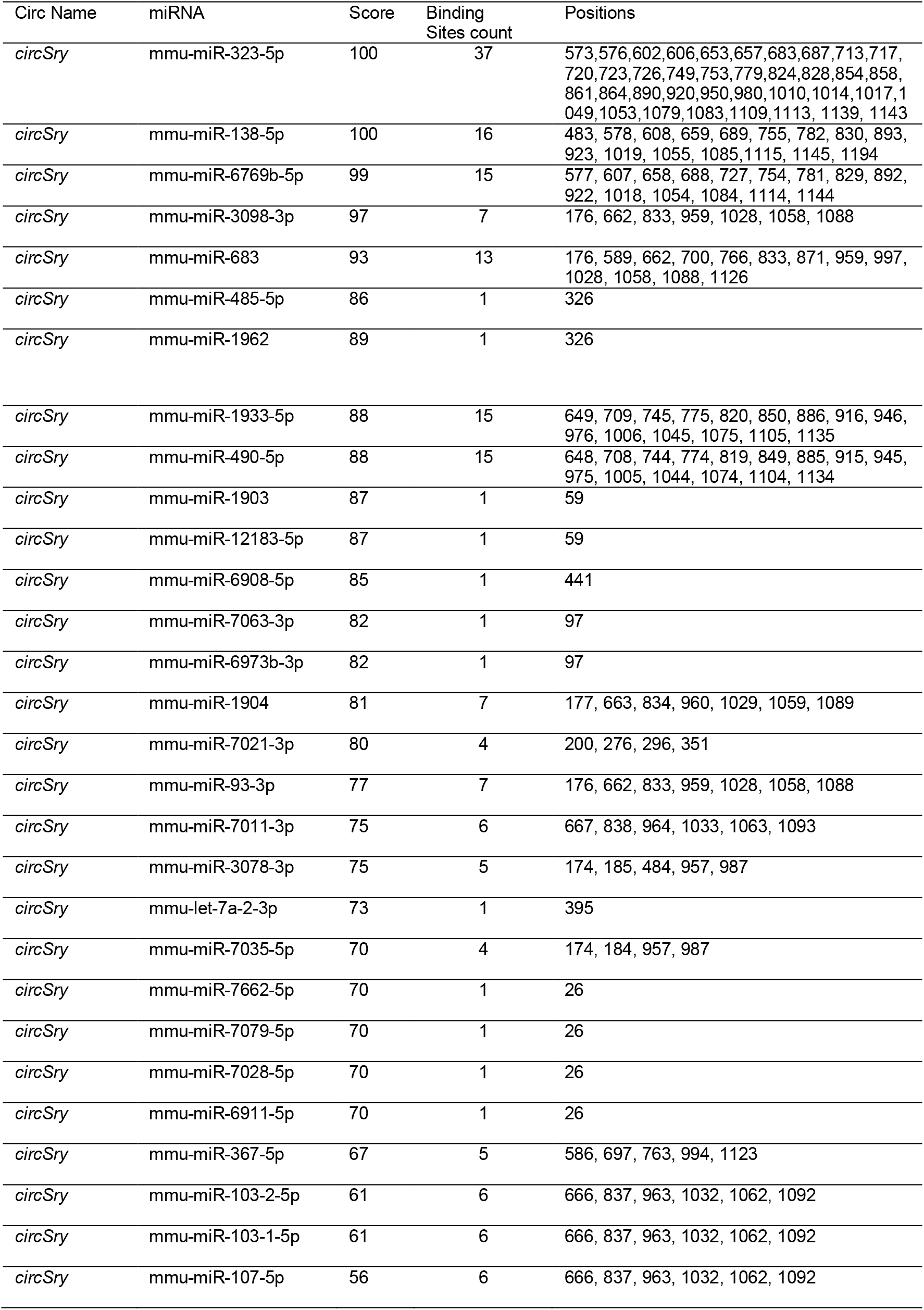
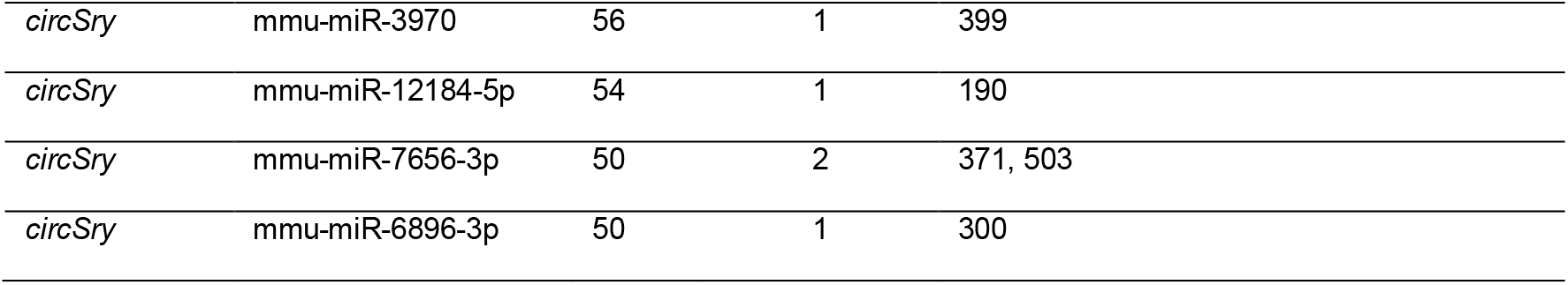
Predicted miRNAs that bind *circSry*

